# Role of cell polarity dynamics and motility in pattern formation due to contact dependent signalling

**DOI:** 10.1101/2020.10.09.331009

**Authors:** Supriya Bajpai, Ranganathan Prabhakar, Raghunath Chelakkot, Mandar M. Inamdar

**Affiliations:** IITB-Monash Research Academy, Mumbai 400076, INDIA; Department of Civil Engineering, Indian Institute of Technology Bombay, Mumbai 400076, INDIA; Department of Mechanical and Aerospace Engineering, Monash University, Clayton, VIC 3800, Australia; Department of Physics, Indian Institute of Technology Bombay, Mumbai 400076, INDIA

## Abstract

A key challenge in biology is to understand how spatiotemporal patterns and structures arise during the development of an organism. An initial aggregate of spatially uniform cells develops and forms the differentiated structures of a fully developed organism. On the one hand, contact-dependent cell-cell signalling is responsible for generating a large number of complex, self-organized, spatial patterns in the distribution of the signalling molecules. On the other hand, the motility of cells coupled with their polarity can independently lead to collective motion patterns that depend on mechanical parameters influencing tissue deformation, such as cellular elasticity, cell-cell adhesion and active forces generated by actin and myosin dynamics. Although modelling efforts have, thus far, treated cell motility and cell-cell signalling separately, experiments in recent years suggest that these processes could be tightly coupled. Hence, in this paper, we study how the dynamics of cell polarity and migration influence the spatiotemporal patterning of signalling molecules. Such signalling interactions can occur only between cells that are in physical contact, either directly at the junctions of adjacent cells or through cellular protrusional contacts. We present a vertex model which accounts for contact-dependent signalling between adjacent cells and between non-adjacent neighbours through long protrusional contacts that occur along the orientation of cell polarization. We observe a rich variety of spatiotemporal patterns of signalling molecules that is influenced by polarity dynamics of the cells, relative strengths of adjacent and non-adjacent signalling interactions, range of polarized interaction, signalling activation threshold, relative time scales of signalling and polarity orientation, and cell motility. Though our results are developed in the context of Delta-Notch signalling, they are sufficiently general and can be extended to other contact dependent morpho-mechanical dynamics.

## I. INTRODUCTION

Morphogenesis is a complex phenomenon in which mechano-chemical pattern formation plays a key role [1]. A large number of self-organising, regular, spatio-temporal patterns in tissues have already been documented. These include, for example, bristle patterns on the *Drosophila notum* [2, 3], spotted skin patterns on pearl danio fish and striped skin patterns on zebrafish [4, 5]. Such spatially differentiated patterns are formed from an aggregate of uniform cells due to cell differentiation process that acquires a different fate depending on their spatial position [6].

Tissues establish self-organizing chemical patterns by interacting chemically and mechanically [7, 8]. Reaction and diffusion processes involving activators and inhibitors can result in a large variety of the so called Turing patterns in the tissues [9, 10]. However, various investigations indicate that a large number of self-organised patterns can also be generated through juxtacrine signalling that occurs through contacts either between cells or between cells and the extra-cellular matrix (ECM) [11]. While Notch signalling is the most prominent juxtacrine developmental signalling pathway, others such *as* Epidermal Growth Factor Receptor (EGFR) and Hedgehog (HH) pathways are also important for morphogenesis [2, 12, 13]. The cellular contacts during signalling could either be local and between the nearest neighours [14–16] or they could be long-ranged and mediated by protrusions such as filopodia [2, 4, 17–19]. Although protrusion-based signalling through Notch pathway is quite common during morphogenesis [2, 18], similar long-ranged signalling is also seen, for example, in Sonic hedgehog (Shh) during limb patterning in vertebrates [17].

In contact based signalling, lateral inhibition is a cell interaction process where a cell with a particular fate inhibits the other cells in contact from achieving the same fate [20]. In embryos and adults, a number of genes and proteins of neurogenic class are involved in lateral inhibition signalling [20–22]. A transcriptional regulator TAZ is also recently identified as a mediator of lateral inhibition in zebrafish and is involved in directing cell fate [23]. More commonly, in several multi-cellular organisms such as flies, worms, fish and other vertebrates, signalling takes place by the lateral inhibition of Delta and Notch, which are trans-membrane molecules that reside on the cell surface [24, 25]. In the rest of the paper, we focus on lateral inhibition by Delta-Notch signalling

The short-range signalling via lateral inhibition, in which the immediate neighbouring cells in a tissue attain a different fate results in a self-organised checker-board pattern [24]. On the other hand, more complex patterns such as bristle patterns in *Drosophila notum* and stripe patterns in zebrafish can be produced by protrusion-mediated long-range signalling with protrusion directionality and signalling efficiency [26, 27]. Moreover, the patterning dynamics can be sped up by the inclusion of the mutual inactivation of Delta-Notch along with the dynamics of lateral inhibition [28, 29].

Coupling between collective cell migration, cell mechanics, and cell-cell signalling is observed in many biological processes such as wound healing, cancer metastasis, branching morphogenesis and embryonic development [30–32]. This coupling is also observed in the case of Delta-Notch signalling. For example, in endothelial cells exhibiting Delta-Notch kinetics, the expression of Dll4 (Delta) is significantly enhanced at the tips of the migrating epithelium during angiogenesis [33]. Also, Delta increase is associated with the motility and spreading of individual keratinocytes [30] and stimulated lamellipodia formation [34]. Furthermore, Delta-induced activation of Notch is linked with the application of mechanical force [35, 36]. Thus there are good indications that spatiotemporal chemical patterns of molecules due to contact-based signalling are associated with cell-cell signalling kinetics, tissue mechanics, cell polarisation dynamics, and cell motility.

The chemical patterns due to contact-based signalling are interpreted using models generally with a simplifying assumption that the tissue morphology is fixed and does not alter during the patterning process [2, 24, 26, 28, 29]. This assumption may not always be correct since cell migration and cell division can dynamically modify the connectivity among cells. Hence, in order to maintain a regular pattern, the signalling pathway requires some feedback mechanisms to coordinate with cell migration and dynamic tissue topology. For example, it is known that FGF and Notch signalling pathways play a crucial role in cell fate decisions and cell migration during gastrulation in *Xenopus* [37]. During somitogenesis in zebrafish, it is observed that Delta-Notch signalling is accompanied by cellular movements in the course of segmentation clock generation [38, 39]. Similarly, it is reported that somitogenesis in chick embryos involves a complex interplay of individual cell movements and dynamic cell rearrangements [40]. Such large scale cellular movements and rearrangements of different types of cells are also observed during germ-layer formation in zebrafish [41, 42].

Computational studies show that cell migration plays a vital role in Delta-Notch patterning in zebrafish [43]. Numerical modelling also shows that during somitogenesis, the synchronization of the segmentation clock is sustained and promoted by randomly moving cells [7], which in turn promotes the flow of information across the tissue by cell mixing and destabilizing the regular patterns [8]. In such a case, the ratio of time scales of cell migration and cell-cell signalling is crucial for patterning and information transfer between the moving cells [43].

As discussed above, the cell movement characteristics can control the signalling patterns in the tissues. However, the migration pattern of cells in the tissue strongly depends on mechanical properties and cell polarisation dynamics [44]. It is known that the motile cells mechanically interact via elastic forces, contractile forces, cell-cell adhesive forces as well as active forces [45, 46]. The tissue shows a transition from solid to fluid behaviour depending on the target shape index, migration and the polarity dynamics the cells [44]. The cells can orient randomly or exhibit polar alignment and move collectively depending on the persistence time of cell tracks and the local orientation order between them [47, 48]. For example, collective motion with velocity or polar alignment between cells shows the presence of highly dynamic, large-scale moving structures which shows a lane-like or band-like movement of cells in a tissue [49]. Hence it is important to understand the connection between signalling patterns and cell polarisation and migration dynamics.

Although some of existing theoretical models investigate the potential mechanisms that could result in a variety of patterns due to contact-based signalling, to the best of our knowledge, there are no theoretical studies yet that attempt to include the role of tissue mechanics, cell polarisation dynamics, and cell motility influencing them. However, as discussed above, these factors are expected to be important in dictating orientation, range and topology of cellular contacts in the tissue and hence could be critical for the origin and maintenance of the chemical patterns. To test the influence of above mentioned factors in the patterns formed by signalling molecules, we study the system using the well-established vertex model [45, 50, 51], with several crucial additions. First, we overlay the lateral inhibition based signalling kinetics to the vertex model. We consider both short-ranged as well as long-ranged signalling kinetics, to account for junctional and protrusional contacts [2, 24, 26]. We also study the effect of the activation threshold for long-range signalling on the chemical patterns. Second, we couple the orientation of protrusional contacts with the underlying cell polarities and study the effect of polarisation dynamics on the generated patterns. We specifically look at two cases of polarity dynamics: (i) random rotational diffusion and (ii) polarity alignment with the nearest neighbors. Finally, in addition to cell-signalling, for every cell, we also include cell motility that is oriented along cell polarity and investigate the role of cell migration and tissue mechanics on the resulting signalling patterns due to dynamically evolving cell-cell contacts.

Based on these new inclusions to the model, in addition to the standard checker-board patterns for signalling molecules, we obtain a large number of intricate patterns ranging from well-defined spotted motifs to diffuse patterns. Moreover, for neighbour aligned polarity dynamics, we see striped patterns of signalling molecules. Upon addition of motility, interestingly, we find that the patterns in signalling molecules are maintained, but the cellular structure keeps dynamically shifting in space. We systematically quantify the spatio-temporal characteristics of the chemical patterns by obtaining the number of clusters, cluster size distribution and cluster anisotropy of the signalling molecules, as well as dynamic correlation function. Overall, we show that the dynamics of cell polarity and cell motility greatly influences the richness of molecular patterns arising from contact-based signalling.

## II. METHODS AND MODEL

In our paper, the mechanics of the tissue is implemented using a vertex model [44–46, 51], in which the tissue is represented as a monolayer formed of polygonal cells having vertices and edges. The mechanical forces within the tissue arise from area elasticity and boundary contractility of individual cells and the forces at cell-cell contacts from acto-myosin contractility and E-cadherin adhesivity. The mechanical contribution from these sources can be expressed using a work function of the form,

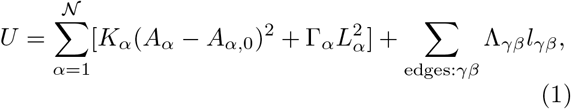

where, is the total number of cells in the monolayer and *K*_*α*_, *A*_*α*_, *A*_*α*,0_, Γ_*α*_, and *L*_*α*_ are the area stiffness, current area, preferred area, boundary contractility and perimeter, respectively, of cell *α*. Λ_*γβ*_ is the contractility of the junction of length *l*_*γβ*_ shared by cells *γ* and *β*. The contributions from these different forcing terms is converted in effective force acting on any vertex *i* as

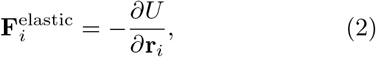

where **r**_*i*_ is the position vector for vertex *i* [50]. In many epithelial tissues, cells are known to be front-rear polarised and in many cases also have self-propelled motility. Hence, in addition to the elastic forces, for a given vertex *i*, we add a motile force [44, 52] of the form

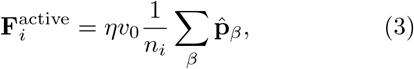

where *n*_*i*_ is the number of cells *β* that contain vertex *i*, 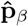_*β*_ is the polarity unit vector for cell *β, η* is the viscous drag acting on the vertex and *v*_0_ is the motility of a single cell. The total force on vertex *i*, which is a combination of the elastic and active force, is balanced by the external viscous force. The resulting dynamical equation of evolution for the vertex position is

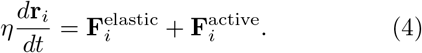

As is common for vertex models, T1 transitions are also included in our formalism and facilitate fluidisation of the tissue.

We model the polarity of every cell to have a tendency to orient with respect to the director (*±* **p**) of its nearest neighbors while also undergoing rotational diffusion [44, 53–56]. This rule can be expressed with the following equation

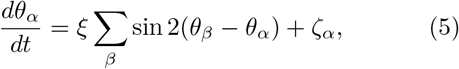

where *θ*_*α*_ denotes the orientation angle of cell polarity, 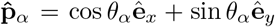. Here, *ξ* is the strength of the polarity alignment of a given cell *α* with respect to that of its connected cells *β* and *ζ*_*α*_ is the rotational noise which follows

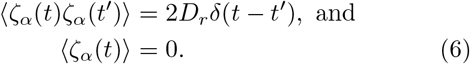

In order to study pattern formation of molecules due to contact-based, cell-cell signalling, we now overlay the signalling kinetics on the mechanical vertex model. As discussed earlier, we use Delta-Notch signalling, which is based on contact based lateral inhibition, as our model system [2, 24, 26]. In our formalism, the Delta-Notch kinetics of the cells is modelled by keeping track of Notch and Delta concentration *N*_*α*_ and *D*_*α*_, respectively, in each cell *α*. It is known that while Notch concentration in a given cell *α* increases with the increase in Delta concentration of the cells in contact, the Delta concentration of that cell decreases with increase in its Notch concentration. This signalling dynamics could mathematically be represented as follows

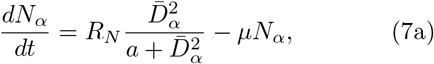

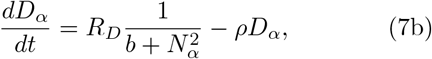

where *R*_*N*_| *µ* and *R*_*D*_| *ρ* are, respectively, the production | decay rates of Notch and Delta. Here, 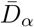 denotes the mean Delta concentration in the cells that are in direct contact with cell *α* through cell-cell junctions and cellular protrusions. More specifically,

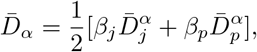

where *β*_*j*_ and *β*_*p*_ correspond to the contact weights for nearest neighbor and protrusional contacts, respectively, such that *β*_*j*_ + *β*_*p*_ = 1. We define

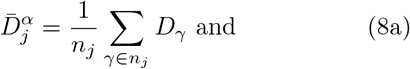

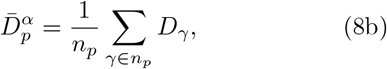

where *n*_*j*_ and *n*_*p*_ are the number of cells in contact with cell *α*, respectively, via cell-cell junctions and protrusions. The nearest neighbors of the cell *α* constitute *n*_*j*_ and the non-adjacent neighbors constitute *n*_*p*_.

There are various readouts for the front-rear polarity of cells such as, the gradient of small GTPase molecules within the cells, gradient in the strength of focal adhesions, and the orientation of golgi with respect to cell nuclei, to name just a few, and signalling pathways such as MAPK are involved in cell polarisation [57, 58]. A free cell typically has motility along the direction of polarity via lamellipodia protrusions at the front. In many cases, filopodial protrusions are also formed at the front and the rear of the cells to aid various aspects of cell migration. Because of this connection between cell polarity, cell motility and filopodial protrusions, in our model we quite reasonably assume that the orientation of cell protrusions is the same as that of cell polarity 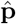 as defined above. Below we outline the procedure followed to obtain the *n*_*p*_ cells that contact cell *α* through protrusions.

The coupling between the mechanical vertex model and the signalling kinetics is made by identifying that the protrusions of cells are indicative of polarity and motility of the cells [59, 60]. In that spirit, cell protrusions are modelled by assuming a protrusional length *l* extending along the orientation of cell polarization, 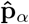 and 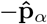 [26]. We assume that the cellular protrusions lie in a interval of [−Δ*θ*,Δ*θ*] around the directions 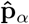 and 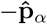 .We choose the protrusion length *l* of cell protrusion [2]. As there is strong experimental evidence of Delta-Notch signalling arising from contact between filopodia of cells [2, 17–19], we focus exclusively on this mode of long range signalling in our paper. Hence, we assume that signalling takes place when the protrusion of a given cell makes contact with the protrusion of other cells within an annulus of thicknessΔ*l* around the protrusion length *l* of the protrusion. Thus effectively, protrusions of two cells can potentially contact with each other for signalling only if the distance between the centers of the cells is within the interval [2*l −Δl*, 2*l*] – let us term this as separation criterion. However, the extent of cellular protrusions overlap depends on the relative positions of cell pairs, polarity of each cell and the angular sweep of protrusions 2Δ*θ* (Fig. 1). For example, ifΔ*θ* = *π/*2, then any pair of cells that satisfy the separation criterion will be in large protrusional overlap with each other. Similarly, ifΔ*θ* is small, then any pair of cells satisfying the separation criterion would have relatively small protrusional overlaps, and that too for only certain relative positions and polarity orientations (Figs. 1bc).

**FIG 1.**
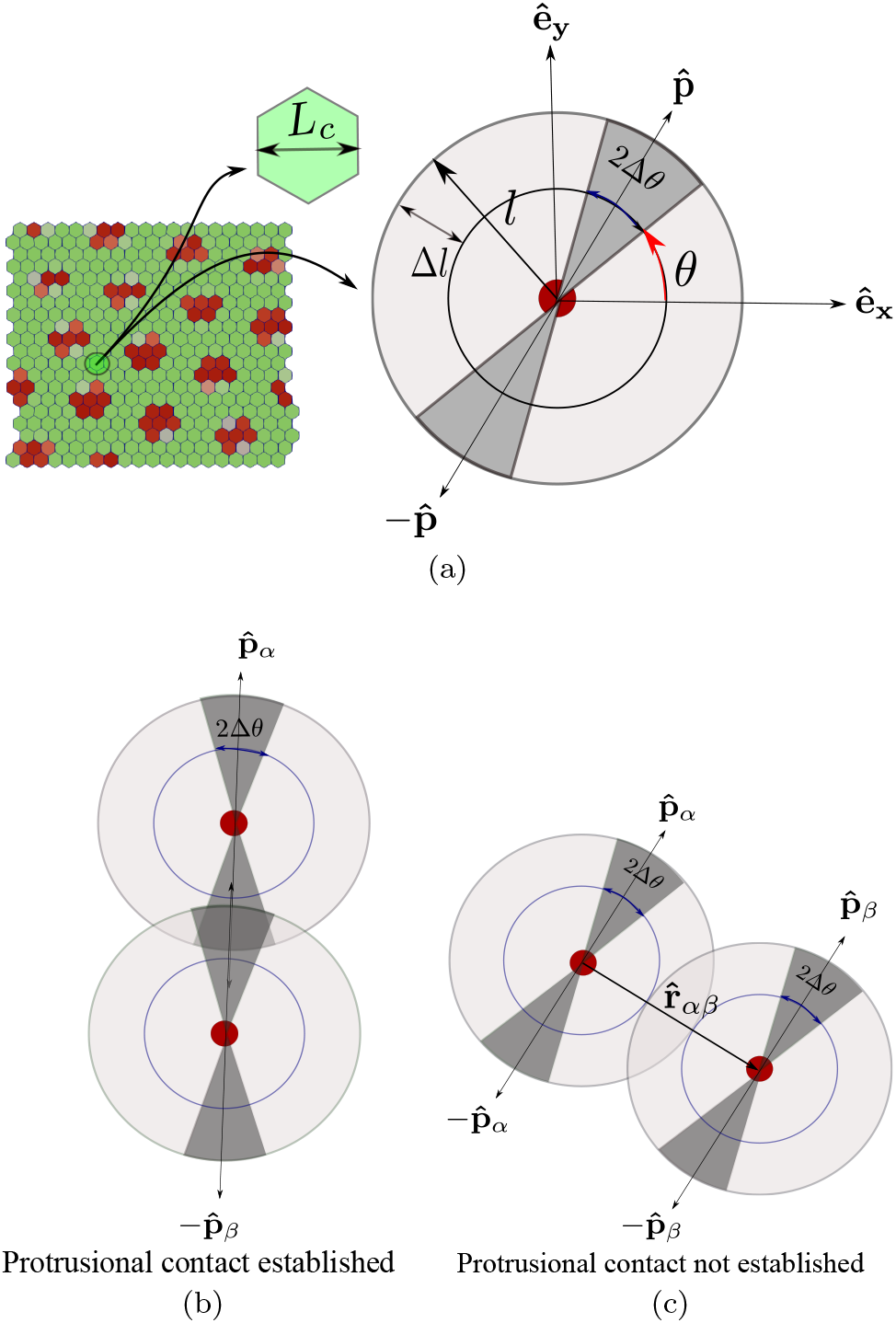
Schematic showing protrusion along cell polarization and contact interactions with neighbors. (a) Single cell with polarisation 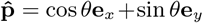and protrusions along 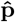 and 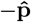. Length of protrusions is *l* and its angular spread is 2Δ*θ* around *θ* and *θ* + *π*. (b) Cellular protrusions of two cells overlapping each other. Likelihood of contact for two cells is high if their cell polarization vectors are coaxial or the angular range of protrusion is high. (c) The centers of two cells that are within range [(2*l −*Δ*l*), 2*l*] of one another but the protrusions do not touch.

The signalling between the contacting cells is believed to have an activation threshold [2] that, for example, could depend on the extent of the overlap [61– 63]. The exact protrusional overlap between the pair of cells can be calculated using geometry. However, since in our model we couple the protrusion orientation with the cell polarity which constantly evolves in time (Eq. 5), for computational convenience we use a simpler criteria for overlap that also includes the signalling activation threshold *T* in a coarse-grained fashion. In our model, we define

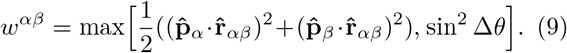

Only if *w*^*αβ*^*≥ T* and the cell separation criterion is satisfied there exists protrusional contact between the cell pair *α, β*. Here, 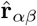 is the unit vector from the center of cell *α* to the center of cell *β* (Fig. 1c). The threshold criteria excludes, in a coarse-grained manner, unfavourable configurations from forming signalling contacts.

The different parameters used in our model are non-dimensionalised as discussed in Table I (Electronic supplementary material).

## III. RESULTS

As described in Section II, there are different factors that interact with each other to control the fate of Delta-Notch pattern formation. Some of these are the relative contributions from junctional and protrusional contacts (*β*_*j*_*/β*_*p*_), Delta-Notch signalling rates (*ρ≈ µ ≈R*_*D*_ *≈R*_*N*_), polarity orientation time-scales (1*/D*_*r*_, 1*/ξ*), length and overlap margin of protrusions (*l,Δl*), angular range of protrusions (Δ*θ*), and neighbour exchange time-scales (*L*_*c*_*/v*_0_), where *L*_*c*_ is the characteristic length scale of the system given by the cell size (Fig. 1a). For example, in the case, whenΔ*θ* is large, the contact between any pair of cells only depends on the spacing between the cells. Hence, the pattern formation is expected to be predominantly dictated by the relative time-scales *L*_*c*_*/v*_0_ over which the cells move away from each other and 1*/ρ*. On the other hand, whenΔ*θ* is small, even if the spacing between the cells does not change (e.g., when *v*_0_ ≈ 0) the pattern formation from protrusional contacts should still be influenced by the time-scales for polarity changes 1*/D*_*r*_ when compared with the signalling time-scales 1*/ρ*. In this section, we systematically explore, how these different chemical and mechanical factors decide the spatio-temporal dynamics of signalling patterns. In Sec. IIIA-C, we study the role of mechanochemical parameters on signalling patterns when cell motility is low. The effect of cell motility on signalling patterns is explicitly investigated in Sec. III D. The mechanical parameters for the vertex model were chosen such that the tissue remained in the solid-like regime (Sec. IIIA-C) or in the fluid-like regime (Sec. IIID) [44, 46]. The dynamical parameters were chosen such that the signalling time-scales allowed for the chemical patterns to form, but the polarity and motility time-scales could also allow the patterns to remain dynamic.

### A. Role of contact ratio (*β*_*j*_*/β*_*p*_) from junctional and protrusion-mediated contacts

The strength of Delta–Notch signalling at junctional and protrusional contacts is captured by *β*_*j*_ and *β*_*p*_, respectively. The contact ratio *β*_*j*_*/β*_*p*_ is critical for deciding signalling pattern in this model. When the contact ratio is large *β*_*j*_*/β*_*p*_ ≫ 1 checker board pattern emerges since the signalling is dominated by the junctional contacts as in the classic model by Collier et al. [24] (Fig. 2a) and (Electronic Supplementary Material Movie 1). On the other hand, consider the case of small contact ratio *β*_*j*_*/β*_*p*_ ≪ 1 with*Δθ* = *π/*2 (Fig. 2b) and (Electronic Supplementary Material Movie 2). Here, the dominant mode of signalling is through cell protrusions. Moreover, for*Δθ* = *π/*2, the cell-cell signalling is isotropic and occurs for any pair of cells that satisfy the separation criterion. The signalling pattern in this case is similar to the checkerboard pattern observed for large contact ratio. However, the pattern shows a new length scale corresponding to the size of protrusions (2*l ≈* 3 cell-lengths) [26]. Thus the nature of pattern formation in contact based signalling is influenced by the relative strengths of junctional and protrusional contacts.

**FIG 2.**
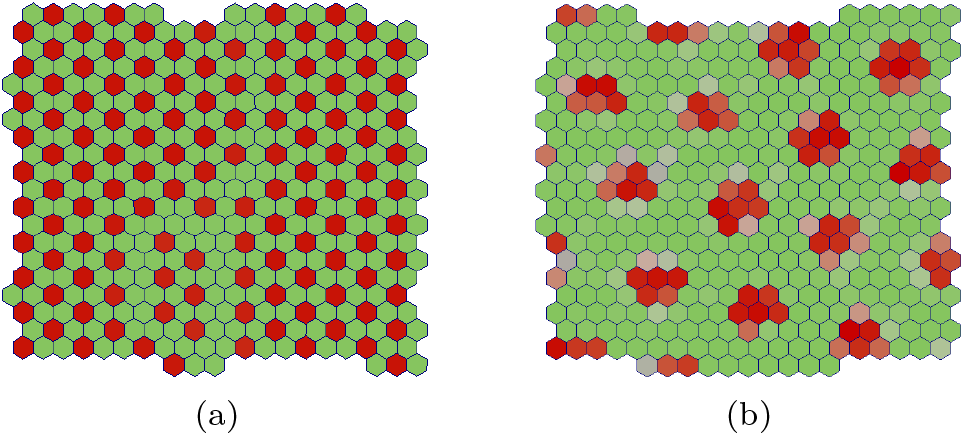
signalling patterns formed by contact mediated signalling via (a) junctional contacts *β*_*j*_ *≫β*_*p*_ and (b) protrusional contacts *β*_*p*_ *≫β*_*j*_. The final time point of the simulations after steady pattern has emerged is shown for both the cases. The results in (a) and (b) confirm the findings in Refs. [24, 26], respectively.

### B. Role of angular range of protrusions (Δ*θ*) and activation threshold (*T*)

As discussed in Sec. II, long-range signalling can be achieved by protrusional contacts. As described there, the orientation of protrusion for any cell *α* is decided by the direction of cell polarization 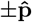 In this section, we consider the case where the cell polarization is governed by the random rotational diffusion only (*ξ* = 0). The signalling dynamics additionally depend on the length and overlap range of protrusion (*l*,Δ*l*), the angular range of the protrusionsΔ*θ* and activation threshold (*T*). We now systematically study the effect ofΔ*θ* and *T* on signalling patterns by varying only these two while keeping all other model parameters fixed (Fig. 3) and (Electronic Supplementary Material Movie 2-7). The protrusions are more polarized ifΔ*θ* is small and not polarized at all ifΔ*θ* = *π/*2, i.e., the protrusions can grow in all directions.

**FIG 3.**
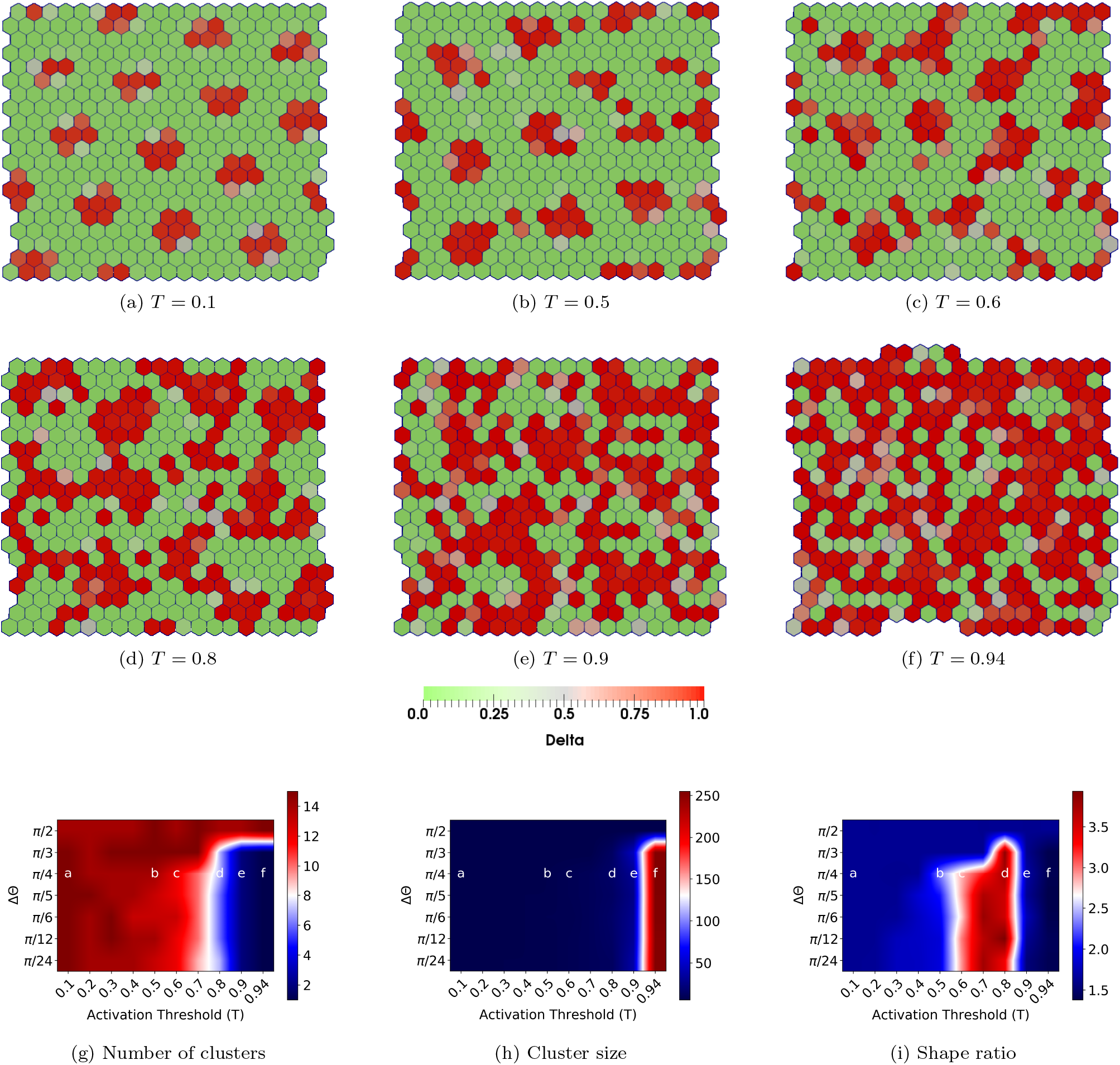
Role of angular range of protrusionsΔ*θ* and activation threshold *T* on pattern formation. (a-f) Steady-state Delta(red)-Notch(green) patterns obtained with *R*_*N*_ = *R*_*D*_ = *ρ* = *µ* = 1, *D*_*r*_ = 10^*−*3^, 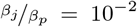,Δ*θ* = *π/*4, *v*_0_= 3.1*×*10^*−*4^, and Λ = *−*13.77, *ξ* = 0 and varying *T ∈* [0.1, 0.5, 0.6, 0.8, 0.9, 0.94]. (g,h) Phase diagrams for the median number of clusters and the median cluster size (median number of Delta cells per cluster) (see Electronic supplementary material Section II) in a confluent tissue as a function ofΔ*θ* and *T*. Large value of cluster number with small cluster size indicates many isolated small Delta patches, whereas a small number of clusters with large cluster size indicates connected regions of Delta expression. (i) Phase diagram for the median shape ratio of Delta clusters in a confluent tissue as a function ofΔ*θ* and activation threshold *T* (see Electronic supplementary material Section III). Lower and higher values of this quantity indicate dominant presence of circular and elongated patches, respectively.

We keepΔ*θ* = *π/*4 and explore how the steady-state signalling patterns evolve with the activation threshold *T*. When *T* is relatively small, we see isolated, ordered patterns of sharp isotropic spots of Delta expression, similar to the ones already discussed in Sec. III A (see Fig. 2b). Upon increase in *T*, there is a decrease in Delta-Notch signalling (Eq. 9) that results in reduction of Notch in cells and hence a general increase in Delta levels (Eq. 9). Moreover, the Delta cell patches also gets relatively anisotropic. As a result, the Delta expression patterns start getting less structured, more elongated, and increasingly connected. This effect becomes most pervasive at the largest threshold value.

We now quantify different aspects of Delta patterns that are observed for various combinations ofΔ*θ* and *T*. To get insights into the connectivity of the Delta patches, we compute the median number of Delta clusters and median size (number of Delta cells per cluster) of isolated Delta clusters. We define one cluster of Delta cells as the group of connected cells, each with Delta concentration *D > D*_critical_ (see Electronic supplementary material Section II). To also get insight into the geometry of these patches, we then quantify their shape ratio (see Electronic supplementary material Section III). In Figs. 3g, 3h and 3i, respectively, we represent the median number of clusters, median size of the clusters and their median shape ratio, calculated over space, time and simulation runs, as functions ofΔ*θ* and *T* (see Electronic supplementary material Section II and III). By observing these phase-diagrams together we can see that, for lower values of *T* the Delta expression patterns are isolated in small isotropic clusters (Figs. 3ab). However, upon increase in *T*, the clusters keep getting smaller in numbers, i.e, larger in size, and become increasingly elongated for lower values ofΔ*θ* (Figs. 3cd). For largest values of *T*, the clusters remain bigger but become more isotropic due to increasing connectivity of Delta regions (Fig. 3ef). For large values ofΔ*θ*, however, the clusters always remain small and isotropic, as discussed in Sec. III A.

We note that, whenΔ*θ* = *π/*4 and *T ≤* 0.5, from Eq. 9, *w*_*αβ*_ *≥ T*, due to which the results in Fig. 3a (*T* = 0.1) and Fig. 3b (*T* = 0.5) should ideally be identical. However, due to small round-off errors during computing, at these values of *T* and Δ*θ* the condition *w*_*αβ*_ *≥ T* is always satisfied for *T* = 0.1 but not for *T* = 0.5. Consequently, there are a few differences between Figs 3a and 3b. This small numerical deviation would be the expected at the critical transition point when sin^2^ Δ*θ* = *T*.

We also point out that, for large values of *T*, the Delta patterns depend on protrusion orientations 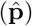, which evolve on time-scales set by 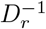. When *D*_*r*_ is zero, we get a static but well-formed Delta pattern (Electronic Supplementary Material Movie-23). When *D*_*r*_ is non-zero but small compared to signalling rates (Eq. 7), we get well-formed but fluctuating Delta patterns (Electronic Supplementary Material Movie-4). However, when *D*_*r*_ becomes large, the protrusional contacts evolve too fast for signalling to take effect, because of which the patterns are underdeveloped as Delta levels remain low and fluctuating (Electronic Supplementary Material Movie 24). Hence, in this section, we chose *D*_*r*_ that was small compared to signalling rates to get well-formed but dynamic Delta patterns (also see Electronic Supplementary Material Section IV).

We thus find that a rich array of Delta-Notch patterns are observed due to an interplay between the angular range of protrusions and the threshold for signalling due to protrusional contacts. We also provide an effective way of quantifying the nature of these patterns.

### C. Role of coupling strength ratio (*ξ/D*_*r*_) on pattern formation

In our model, the dynamics of cell polarity has two components (Eq. 5). The first component tends to align the polarity of any cell with that of its nearest neighbors with rate *ξ* and attempts to bring about global alignment of polarity in the tissue [49]. The second component *D*_*r*_ brings about rotational diffusion of cell polarity, thus creating an overall disorder in tissue polarity. As studied in the previous section, for the case of polarity alignment rate *ξ* = 0, the cell polarities dictate the local dynamics of protrusional contacts (Eq. 9) and hence the Delta-Notch patterns. However, since *ξ* influences the global alignment of cell polarity, in this section we study the role of the coupling strength ratio *ξ/D*_*r*_ on Delta-Notch pattern formation.

We fix *T* = 0.5, *D*_*r*_ = 0.1, Δ*θ* = *π/*4 and vary the value of *ξ* from 0*−* 0.25. The progressively changing patterns for increasing magnitude of *ξ/D*_*r*_ are shown in (Figs. 4a-f) and (Electronic Supplementary Material Movies 8-13). As expected, when *ξ* is relatively small, *D*_*r*_ dominates and the cell polarity is spatially disordered, thus resulting in isotropic circular patterns of Delta expression (also see Fig. 3a). However, when *ξ* becomes comparable to *D*_*r*_, the spatial disorder of cell polarity decreases and local regions of polarity alignment with an effective direction are created. Since protrusions are oriented along the polarity of cells in our model, the cell-cell contacts predominantly occur along the effective polarity orientation and very little in the perpendicular direction. As a result, the signalling is diminished along the perpendicular direction resulting in reduction of Notch levels along that direction. The lowering of Notch concentration, in turn, results in greater expression of Delta, thus leading to formation of elongated Delta domains. Consequently, we see Delta expression emerging in stripe-like patterns that are oriented perpendicular to the overall direction of cell polarity in the ordered region. In the region with disordered polarity, we still observe circular regions of Delta expression (Figs. 4c and 4f). As expected, the thickness of stripes and the diameter of the circular spots are roughly equal to twice the protrusion length (2*l ≈* 3 cell lengths). Upon further increase in *ξ*, the cell polarities align globally, thus resulting exclusively in stripe-like patterns of Delta expression. However, the presence of *D*_*r*_ leads to modification of the global polarity alignment causing the patterns to reorient over longer time-scales (see Electronic Supplementary Material Movies 8-13).

**FIG 4.**
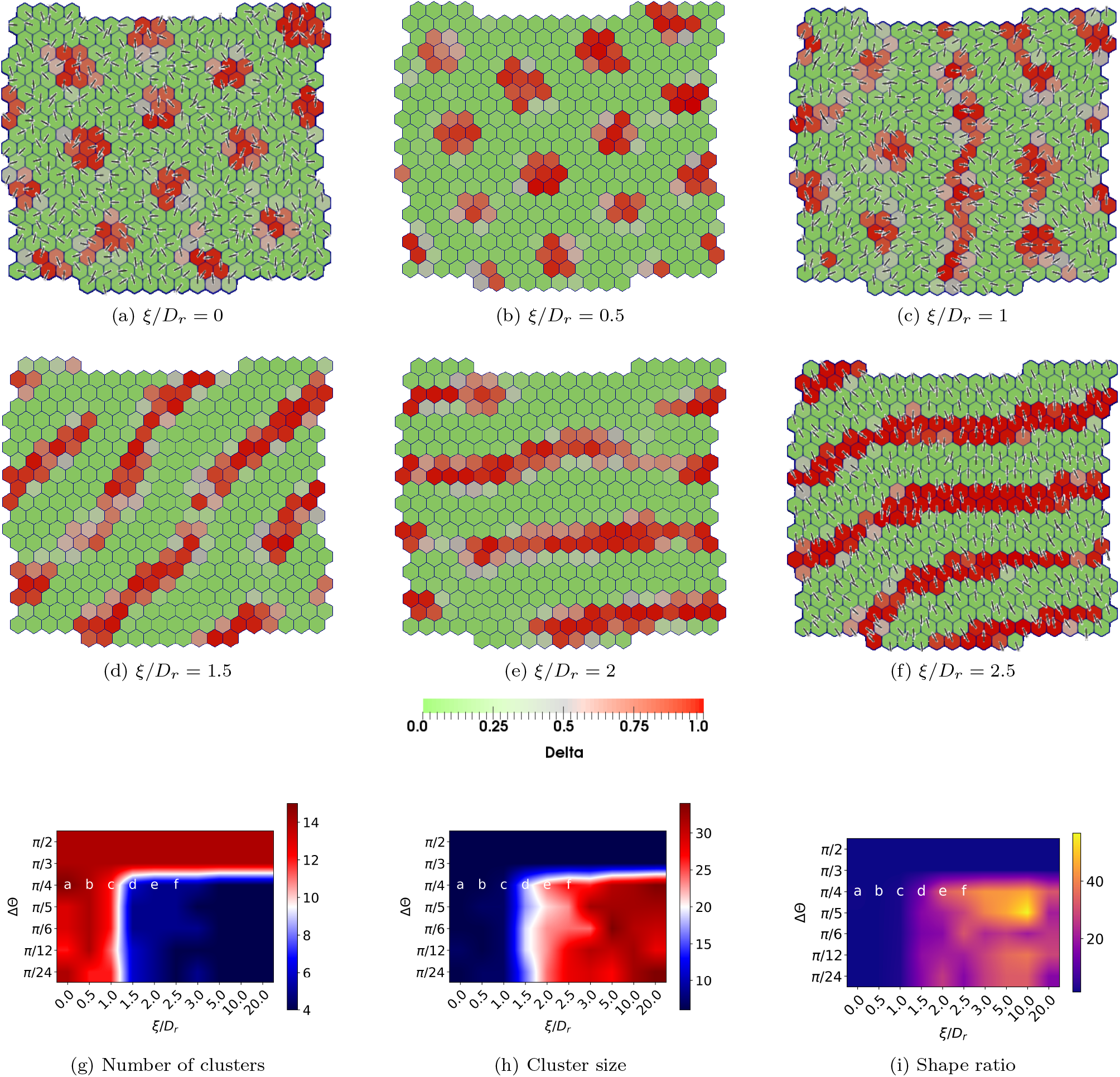
Screenshots showing the steady state Delta(red)-Notch(green) patterns formed with polarized cells with varying coupling strength ratio (*ξ/D*_*r*_) and angular range of protrusions (Δ*θ*). The fixed parameters are *R*_*N*_ = *R*_*D*_ = *ρ* = *µ* = 1, Λ = *−*13.77, *D*_*r*_ = 0.1, *v*_0_ = 3.1 *×* 10^*−*4^, 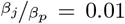, *T* = 0.5, and Δ*θ* = *π/*4. The patterns (a-f) are obtained by varying coupling strength ratio *ξ/D*_*r*_ *∈*[0, 0.5, 1.0, 1.5, 2.0, 2.5]. The lines in each cell indicate the nematic orientation of cell polarity. No arrows are shown due to the equivalence of 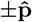 in our model for cell motility (Eq. 5) and protrusional signalling (Eq. 8). (g,h) Phase diagrams for the median number of clusters and median cluster size in a confluent tissue (see Electronic supplementary material Section II) as a function Δ*θ* and *ξ/D*_*r*_. (h) Phase diagram for median shape ratio in a confluent tissue as a function of Δ*θ* and *ξ/D*_*r*_ (see Electronic supplementary material Section III). Large number of Delta clusters with small cluster size and low shape ratio indicate the dominance of isolated circular patterns, wheres low number of clusters with big cluster size and high shape ratio point towards stripe-like patterns.

As we had done in the previous section, we now quantify the median number of clusters, median cluster size and median shape-ratio of the Delta patterns using phase-diagrams obtained as a function of Δ*θ* and *ξ/D*_*r*_ (Figs. 4g-i, also see Electronic supplementary material Section II and III). As expected, for large values of Δ*θ*, we mostly observe patterns of isolated, circular clusters, irrespective of the magnitude of *ξ/D*_*r*_, since the cell protrusional contacts are mostly isotropic. However, for lower values of Δ*θ*, an increase in *ξ/D*_*r*_, which results in cell polarity ordering, leads to the formation of uniformly oriented and continuous Delta stripes.

We also performed simulations for parameters used in Fig. 4 (see Electronic Supplementary Material Movie 16-22) when *T* = 0.94. Since both *T* and *D*_*r*_ are large, the amount of switching of cell contacts due to fluctuations is naturally very high. As a result, the patterns are either not formed or are more dynamic (see Electronic Supplementary Material Movies 16-18). It could be observed that only when the alignment term *ξ* dominates over *D*_*r*_, the fluctuations in polarity and hence in protrusion alignment are reduced. The system is then able to generate and maintain Delta patterns (Electronic Supplementary Material Movies 19-22).

We thus find that polarity dynamics can have a strong influence on the nature of Delta-Notch signalling patterns.

### D. Effect of motility on pattern formation

So far we have studied the role of protrusion spread, signalling threshold and polarity dynamics on the formation of Delta-Notch patterns in tissues. In our model, the polarity dynamics influences the signalling via modification of protrusional contacts. However, as discussed earlier, cell polarity is also connected with cell migration, which in conjunction with cell shape index can control tissue fluidisation through cell neighbor exchanges and thus influence the signalling pattern. Hence, we provide cells with larger values of motility *v*_0_ and adjust cell line tension Λ such that the cell-shape index 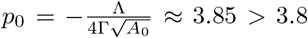, that is required for tissue fluidisation for *v*_0_ = 0 [44, 46, 64]. First, we study the effect of uncorrelated cell movement on pattern formation by fixing *ξ* = 0, *v*_0_ = 0.31, and *D*_*r*_ = 0.001. If the signalling rates are small, then the neighbor exchanges between the cells are too fast as compared to the signalling time-scales. As a result, we do not observe Delta-Notch patterns as for the static cell network. However, upon increasing the signalling rates by ten-fold, we recover back the circular, isolated patterns seen earlier.

Interestingly, the patterns are now no longer static but keep spatially rearranging. The movement of the Delta patterns mainly depends on the dynamics of the cluster of Delta expressing cells, which in turn is dictated by the collective cell migration patterns that are governed by the underlying tissue mechanics and polarity dynamics of individual cells (Figs. 5a-c)(Electronic Supplementary Material Movies 14 and 15). In the case where a particular cluster of Delta cells breaks apart, a new group of Delta expressing cells is created by the entry of new cells into a pre-exisiting nuclei of Delta expressing cells. On the other hand, there are cases where the entire group of Delta expressing cells migrates as a whole in which case the Delta patterns also take the same trajectory as the complete cluster. A combination of cellular movements and chemical patterns leads to an emergent time-scale for the spatial rearrangement of Delta clusters.

**FIG 5.**
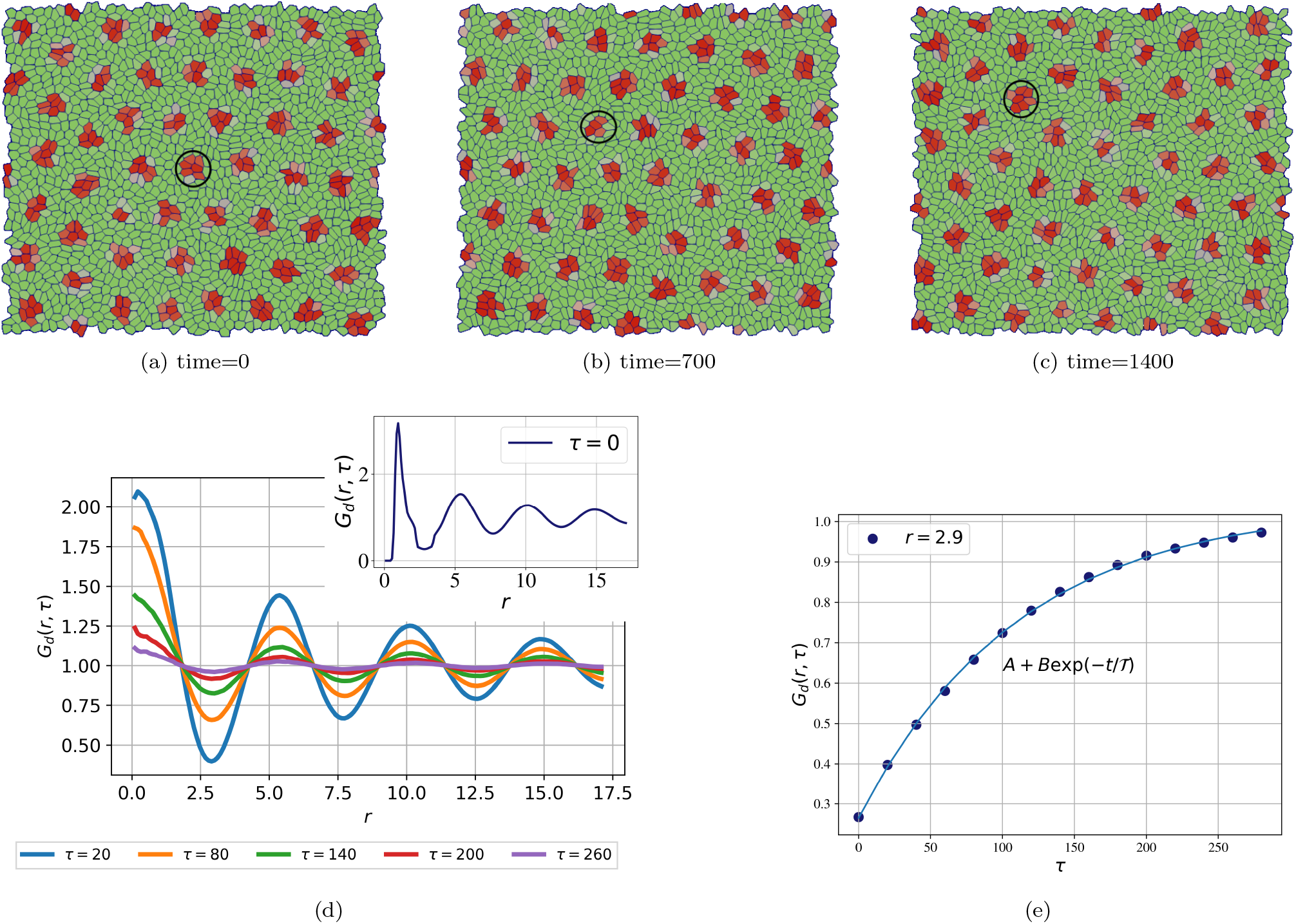
Screenshots and plots showing the effect of cell motility and tissue mechanics on Delta-Notch pattern formation. The parameters used for the simulations are *R*_*N*_ = *R*_*D*_ = *ρ* = *µ* = 10, Λ = *−*14.32, *D*_*r*_ = 0.001, *ξ* = 0, Δ*θ* = *π/*2, *T* = 0.1 and *v*_0_ = 0.31. The shape parameter for the cells *p*_0_ *>* 3.82, the so called fluidisation threshold. (a)-(c) The spot-like Delta patterns keep re-arranging in space as a function of time. The circles correspond to manual tracking of the cluster shown in panels. (d) Plot of the Delta-Delta correlation function *G*_*d*_(*r, τ*) shows clear spatial pattern with a length scale of approximately 5 cell lengths. Although the shape of the function *G*_*d*_(*r, τ*) does not change with *τ*, its amplitude decreases, thus indicating that the dynamic nature of the patterns. (e) The magnitude of *G*(*r, τ*) as a function of *τ* for *r ≈* 2.9 is plotted as a function of time. An exponentially saturating function of the form *A* + *B* exp(*−t/*𝒯) fits well to these values with 𝒯 ≈ 100 and provides the time-scale for pattern re-arrangement.

To quantify the spatio-temporal dynamics of these patterns, we calculate the Delta-Delta radial distribution function, *G*_*d*_(*r, τ*) that is given as

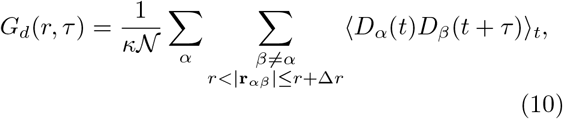

where, *κ* is the normalization factor

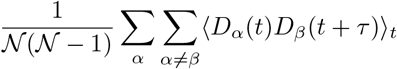

The basic idea behind this function is to capture for every cell *α* at a given time *t* how much does its Delta expression correlate with the Delta levels of every other cell *β* that is present within a particular distance *r* |*<* **r**_*β*_ *−***r**_*α*_| *≤ r* + Δ*r* at time *t* + *τ*. The plots of *G*(*r, τ*) as a function of *r* for different time-lags *τ* are shown in Fig. 5d. The plot for each value of *τ* was obtained from the average of *G*_*d*_(*r, τ*) over three set of simulations, each with 1600 cells. The initial configuration of polarity 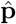 for individual cells was generated from uniform random orientation in the range [*−π, π*] and uniform random concentration of Delta and Notch in the range (0, 1) for a given combination of simulation parameters. When *τ* = 0, we see a decaying oscillatory pattern in space that is indicative of periodic Delta expression with the distance between the centers of neighboring Delta region of approximate 5 cells. For increasing values of *τ*, we see that the shape of *G*_*d*_(*r, τ*) remains invariant, but the amplitude of the function decreases, thus indicating that the patterns are not stationary but diffuse in space. To quantify the rearrangement time-scale of the Delta patterns, we plot the amplitudes of the radial distribution function *G*_*d*_(*r, τ*) corresponding to its first minima *r ≈* 2.5 as a function of time-lag *τ* (Fig. 5e). After fitting an expression of the form *A* + *B* exp(*− τ/ 𝒯*), the pattern re-arrangement time-scale *T ≈* 90 emerges. We note that 𝒯 is much greater than either the time-scale for signalling, (*τ*_*s*_ = 1*/µ ≈* 0.1) or that for cellular rearrangements (*τ*_*r*_ = *L*_*c*_*/v*_0_ *≈* 3) and is very likely an emergent time-scale. Such emergent features are not uncommon in active systems and require a more detailed analysis that is beyond the scope of the current work [65–67].

We also study the effect of cell movement on pattern formation when the alignment strength ratio *ξ/D*_*r*_ is relatively high with *ξ* = 0.25, *v*_0_ = 0.31, and *D*_*r*_ = 0.1. In this case, we see that stripe-like patterns of Delta expression are seen similar to the case when *v*_0_ = 10^*−*4^ (Electronic Supplementary Material Movies 14 and 15). However, the patterns are more dynamic and, as opposed to the formation and breaking of clusters in the case of spot-like patterns when *ξ* = 0, we see that the stripes break and merge to continuously change their alignment.

Thus, we observe that the polarity and motility dynamics of cells, along with tissue mechanics, influence the signalling patterns and hence the spatio-temporal levels of Delta and Notch expression.

## IV. DISCUSSION

In this study, we report a rich variety of Delta-Notch patterns that depend on the nature of cell-cell contacts, signalling threshold, polarity dynamics, cell motility and tissue mechanics. The classic model by Collier et al. [24] exhibits checkerboard pattern for Delta-Notch expression. We show that this pattern modifies to a spot-like like pattern due to long-range contacts with essentially a change in the length-scale that arises due to linear protrusion range. However, the modification in the angular range of protrusional contacts elicits local contact anisotropy and hence results in more elongated Delta patterns. We further showed that the signalling threshold is also important in dictating the connectivity of Delta clusters. Moreover, we systematically quantified the nature of these patterns by measuring the number of Delta cells cluster, cluster size and calculating the shape of individual clusters. We see that by changing the polarity dynamics by increasing the signalling ratio *ξ/D*_*r*_, the cell directors (*±* **p**) become globally aligned thus leading to the formation of stripe-like patterns. We also observed that when the cells have motility and shape index beyond the fluidisation threshold, the cells can rapidly change their connectivity due to which their signalling contacts are also modified. As a result, the expression patterns for Delta-Notch no longer remain static. Their dynamics is decided by the dynamics of the formation and breaking of Delta clusters, which in turn are governed by the motility patterns of the cells. When the polarity diffusion dominates, we see the formation of moving spot-like patterns, which we systematically quantified using the spatio-temporal radial correlation function for Delta expression. On the other hand, when the polarity alignment term dominates, we saw that stripe-like patterns arise. However, unlike for the static case, the stripes keep modifying their alignment by splitting and then merging with the other stripes – this dynamics being governed by cellular movements.

Lateral inhibition is one of the most ubiquitous mode of signalling, and Delta-Notch signalling is the most prominent example of this mechanism. Experimentally, the Delta-Notch signalling mechanism has been studied in detail, and its involvement in cell migration, polarity dynamics, and mechanical aspects of morphogenesis is known. Although, there are a few theoretical models that study the Delta-Notch pattern formation in tissues, there are no theoretical studies on how these patterns are themselves influenced by collective cell dynamics. On the other hand, there are a large number of theoretical studies on collective cell migration, especially on the role of cell motility, polarity and cell shape index on tissue unjamming. However, these studies generally do not consider the effect of tissue kinematics on the underlying signalling patterns. In this study, we combined both these aspects and showed how cell level interactions can lead to tissue level formation of a large variety of Delta-Notch patterns. Although in our model, the Delta-Notch pattern is influenced by cellular dynamics, the signalling itself does not influence the cell dynamics. A next step, for example, would be to include the influence on Delta-Notch levels in the cells on motility and cell-cell adhesivity. We finally note that, although our modeling is developed in the context of Delta-Notch signalling, it is sufficiently general, and provides a broad framework to study the role of collective cell dynamics on chemical pattern formation for any contact based signalling.

## Supporting information

Electronic Supplementary Material

Electronic Supplementary Material - MOVIE CAPTIONS

## ACKNOWLEDGEMENTS

MMI and RC acknowledge Industrial Research and Consultancy Centre (IRCC) at IIT Bombay, India, for financial support. RC thanks Science and Engineering Research Board (SERB), India (Project No. ECR/2017/000744, and SB/S2/RJN-051/2015) for financial support. We thank computing facilities at MonARCH cluster, Monash University.

