## Supplementary material for "Role of cell polarity dynamics and motility in pattern formation due to contact dependent signalling": Electronic Supplementary Material


### I. MODEL PARAMETERS AND NON-DIMENSIONALIZATION

The mechanical energy function and the signalling kinetics equations are non-dimensionalized with characteristic time scale $\frac{\eta}{\Gamma} = 1$ and characteristic length scale $L_c = 1$ (Fig.1a) (Table I).

We simulate a monolayer of tissue with periodic boundary and $\mathcal{N}$ number of cells (no cell divisions or apoptosis). The model is implemented in CHASTE [1] using the C++ libraries. The equation of motion is solved numerically using a simple forward Euler discretization. The signalling equations are solved using Runge-Kutta-Merson method. We choose the time step size $\Delta t = 0.01$ (sufficiently small) to maintain the numerical stability. The initial levels of Notch and Delta concentration are chosen randomly from uniform random number in (0,1) for each cell $\alpha$.

### II. NUMBER OF CLUSTERS AND CLUSTER SIZE

The median number of clusters and the median cluster size is calculated for 400 cells using the density-based spatial clustering (DBSCAN) algorithm [2]. The cell $\alpha$ is considered a Delta cell if the concentration of Delta molecule in the cell is greater than $D_{\text{critical}}$. A group of Delta cells is considered to be in dense region if minimum number of Delta cells in the cluster is 3. Two cells are considered to be touching each other if the Euclidean distance between both the cells are less than or equal to 1.5.

Each data point shown in Figs.3g-i is obtained using five sets of simulations. In each set, for a given combination of parameters, initial polarity $\hat{\mathbf{p}}$ for individual cells was generated from uniform random orientation in the range $[-\pi, \pi]$. Similarly, the initial concentration of Delta and Notch for individual cells was generated from uniform random distribution in the range $(0, 1)$. Then, after removing the transient part of the corresponding simulation, the quantity $X_t^\beta$ (cluster number, cluster size, shape ratio) for each simulation $\beta$ and at every sampling time $t$ was pooled together and its median $\bar{X}$ over $t$ and $\beta$ was used as one data point. The sampling interval for every simulation was $\Delta T = 1$. A similar procedure was followed to obtain Figs.4g-i, except that the total number of simulation runs in this case was six instead of five.


*
†
‡

TABLE I. The model parameters relative to characteristic length scale $L_c = 1$ (Fig.1a) and characteristic time scale $T_c = 1$

| **Dimensionless parameters** | **Parameter values** |
|---|---|
| $L_c$ | 1.0 |
| $\eta$ | 1.0 |
| $\Gamma$ | 1.0 |
| $K \equiv \frac{KT_cL_c^2}{\eta}$ | 1.15 |
| $\Lambda \equiv \frac{\Lambda T_c}{4L_c\eta}$ | $[-13.77, -14.32]$ |
| $v_0 \equiv {}^{v_0T_c}/{}_{L_c\eta}$ | $3.1 \times 10^{-4}, 0.31$ |
| $A_0 \equiv A_0/L_c^2$ | 0.866 |
| $\xi \equiv \xi T_c$ | $0 - 2$ |
| $D_r \equiv D_rT_c$ | $[0.001, 0.1]$ |
| $D_\alpha \equiv {}^{D_\alpha}/{}_{D_0}$ | $0 - 1$ |
| $N_\alpha \equiv {}^{N_\alpha}/{}_{N_0}$ | $0 - 1$ |
| $R_D \equiv R_DT_c$ | $1, 10$ |
| $R_N \equiv R_NT_c$ | $1, 10$ |
| $\rho \equiv \rho T_c$ | $1, 10$ |
| $\mu \equiv \mu T_c$ | $1, 10$ |
| $l \equiv l/L_c$ | 1.7 |
| $\Delta l \equiv \Delta l/L_c$ | 1.2 |
| $\Delta\theta \equiv \Delta\theta$ | $\pi/24 - \pi/2$ |
| $\Delta t \equiv \Delta t/T_c$ | 0.01 |
| $a$ | $[0.01]$ |
| $b$ | $[100]$ |
| $D_{\text{critical}} \equiv {}^{D_{\text{critical}}}/{}_{D_0}$ | 0.5 |
| $\mathcal{N}$ | $[400, 1600]$ |
| $\Delta r$ | 0.1 |

### III. SHAPE RATIO

The shape ratio is calculated for 400 cells. The cell $\alpha$ is considered a Delta cell if the concentration of Delta molecule in the cell is greater than $D_{\text{critical}}$. The inertia matrix of a single cluster is computed as follows:

$$\mathbf{A} = \begin{bmatrix} \mathbf{I}_{xx} & \mathbf{I}_{xy} \\ \mathbf{I}_{xy} & \mathbf{I}_{yy} \end{bmatrix}$$

$$\mathbf{I}_{xx} = \sum_{i=1}^{N_c} (A_i(\mathbf{x}_i - \mathbf{x}_{\text{mean}})^2), \tag{1}$$

$$\mathbf{I}_{yy} = \sum_{i=1}^{N_c} (A_i(\mathbf{y}_i - \mathbf{y}_{\text{mean}})^2), \tag{2}$$

$$\mathbf{I}_{xy} = \sum_{i=1}^{N_c} (A_i(\mathbf{x}_i - \mathbf{x}_{\text{mean}})(\mathbf{y}_i - \mathbf{y}_{\text{mean}})) \tag{3}$$

where, $N_c$ is the number of cells in a cluster. Eigen values of $\mathbf{A}$ is calculated and shape ratio is estimated as the ratio of the maximum and minimum eigen values. The median shape ratio is calculated using the same procedure as described in Section II above.

The median of the shape ratios of all the clusters of a time frame is calculated, and the median of all the shape ratios obtained from all time frames is the final shape ratio.

### IV. ROLE OF $D_r$ AND $T$ IN DELTA PATTERN FORMATION

At lower values of threshold $T$, the differences in the patterns formed by static and rotationally diffusing filopodia are negligible since, in this case, the protrusional contacts between the nearby cells are mostly independent of filopodia orientation ($\hat{\mathbf{p}}$) for intermediate to high values of $\Delta\theta$ (Eq. 9 of the main paper). However, for higher values of $T$, the nature of Delta patterns depend on filopodia orientations and the relative positions of cells (Eq. 9). In this case, when $D_r$ is zero, the final patterns are static and seem to depend only on the initial orientations of filopodia (Movie 23, for $D_r = 0$, $\Delta\theta = \pi/4$, $T = 0.6$, and initial orientation randomly chosen from $[-\pi, \pi]$). When $D_r$ is non-zero but relatively small compared to the Delta-Notch signalling rates (Eq. 7), the system remains established in a particular configuration of orientation network of protrusional connections for a sufficient duration of time to establish Delta levels and patterns. However, due to the non-zero value of $D_r$, the orientations evolve and hence form newer protrusional connection network, thus leading to dynamical patterns (Movie 4, for $D_r = 10^{-3}$, $T = 0.6$ and $\Delta\theta = \pi/4$). Finally, when $D_r$ becomes large when compared to the signalling rates, the protrusional contact network evolves much faster due to which the system does not get sufficient time to establish Delta levels and patterns. Consequently, the overall Delta levels remain low and time-varying, and the patterns not as anisotropic as for the case $D_r = 0$ (Movie 24, for $D_r = 1$, $T = 0.6$ and $\Delta\theta = \pi/4$). Thus, Delta patterns depend on threshold $T$, $\Delta\theta$, and the relative time-scales for orientational changes and signalling.

### V. MOVIE CAPTIONS

**Movie-1** corresponding to Fig.2a. signalling pattern formed by contact mediated signalling via junctional contacts $\frac{\beta_j}{\beta_p} = 99$.

**Movie-2** corresponding to Fig.3a. Pattern obtained using the model for $R_N = R_D = \rho = \mu = 1$, $D_r = 10^{-3}$, $\frac{\beta_j}{\beta_p} = 10^{-2}$, $\Delta\theta = \pi/4$, $v_0 = 3.1 \times 10^{-4}$, and $\Lambda = -13.77$ and $T = 0.1$.

**Movie-3** corresponding to Fig.3b. Pattern obtained using the model for $R_N = R_D = \rho = \mu = 1$, $D_r = 10^{-3}$, $\frac{\beta_j}{\beta_p} = 10^{-2}$, $\Delta\theta = \pi/4$, $v_0 = 3.1 \times 10^{-4}$, and $\Lambda = -13.77$ and $T = 0.5$.

**Movie-4** corresponding to Fig.3c. Pattern obtained using the model for $R_N = R_D = \rho = \mu = 1$, $D_r = 10^{-3}$, $\frac{\beta_j}{\beta_p} = 10^{-2}$, $\Delta\theta = \pi/4$, $v_0 = 3.1 \times 10^{-4}$, and $\Lambda = -13.77$ and $T = 0.6$.

**Movie-5** corresponding to Fig.3d. Pattern obtained using the model for $R_N = R_D = \rho = \mu = 1$, $D_r = 10^{-3}$, $\frac{\beta_j}{\beta_p} = 10^{-2}$, $\Delta\theta = \pi/4$, $v_0 = 3.1 \times 10^{-4}$, and $\Lambda = -13.77$ and $T = 0.8$.

**Movie-6** corresponding to Fig.3e. Pattern obtained using the model for $R_N = R_D = \rho = \mu = 1$, $D_r = 10^{-3}$, $\frac{\beta_j}{\beta_p} = 10^{-2}$, $\Delta\theta = \pi/4$, $v_0 = 3.1 \times 10^{-4}$, and $\Lambda = -13.77$ and $T = 0.9$.

**Movie-7** corresponding to Fig.3f. Pattern obtained using the model for $R_N = R_D = \rho = \mu = 1$, $D_r = 10^{-3}$, $\frac{\beta_j}{\beta_p} = 10^{-2}$, $\Delta\theta = \pi/4$, $v_0 = 3.1 \times 10^{-4}$, and $\Lambda = -13.77$ and $T = 0.94$.

**Movie-8** corresponding to Fig.4a. Pattern obtained using the model with parameter $R_N = R_D = \rho = \mu = 1$, $\Lambda = -13.77$, $D_r = 0.1$, $v_0 = 3.1 \times 10^{-4}$, $\frac{\beta_j}{\beta_p} = 0.01$, $T = 0.5$ and $\Delta\theta = \pi/4$ and $\xi/D_r = 0$.

**Movie-9** corresponding to Fig.4b. Pattern obtained using the model with parameter $R_N = R_D = \rho = \mu = 1$, $\Lambda = -13.77$, $D_r = 0.1$, $v_0 = 3.1 \times 10^{-4}$, $\frac{\beta_j}{\beta_p} = 0.01$, $T = 0.5$ and $\Delta\theta = \pi/4$ and $\xi/D_r = 0.5$.

**Movie-10** corresponding to Fig.4c. Pattern obtained using the model with parameter $R_N = R_D = \rho = \mu = 1$, $\Lambda = -13.77$, $D_r = 0.1$, $v_0 = 3.1 \times 10^{-4}$, $\frac{\beta_j}{\beta_p} = 0.01$, $T = 0.5$ and $\Delta\theta = \pi/4$ and $\xi/D_r = 1.0$.

**Movie-11** corresponding to Fig.4d. Pattern obtained using the model with parameter $R_N = R_D = \rho = \mu = 1$, $\Lambda = -13.77$, $D_r = 0.1$, $v_0 = 3.1 \times 10^{-4}$, $\frac{\beta_j}{\beta_p} = 0.01$, $T = 0.5$ and $\Delta\theta = \pi/4$ and $\xi/D_r = 1.5$.

**Movie-12** corresponding to Fig.4e. Pattern obtained using the model with parameter $R_N = R_D = \rho = \mu = 1$, $\Lambda = -13.77$, $D_r = 0.1$, $v_0 = 3.1 \times 10^{-4}$, $\frac{\beta_j}{\beta_p} = 0.01$, $T = 0.5$ and $\Delta\theta = \pi/4$ and $\xi/D_r = 2.0$.

**Movie-13** corresponding to Fig.4f. Pattern obtained using the model with parameter $R_N = R_D = \rho = \mu = 1$, $\Lambda = -13.77$, $D_r = 0.1$, $v_0 = 3.1 \times 10^{-4}$, $\frac{\beta_j}{\beta_p} = 0.01$, $T = 0.5$ and $\Delta\theta = \pi/4$ and $\xi/D_r = 2.5$.

**Movie-14** corresponding to Fig.5a-c. The parameter values used for the simulations are $R_N = R_D = \rho = \mu = 10$, $\frac{\beta_j}{\beta_p} = 0.01$, $\Lambda = -14.32$, $D_r = 0.001$, $\xi = 0$, $\Delta\theta = \pi/2$, $T = 0.1$ and $v_0 = 0.31$. The shape parameter for the cells $p_0 > 3.82$.

**Movie-15** for the tissue in fluid region with stripe-like pattern. The parameter values used for the simulations are $R_N = R_D = \rho = \mu = 10$, $\frac{\beta_j}{\beta_p} = 0.01$, $\Lambda = -14.32$, $D_r = 0.1$, $\xi = 0.25$, $\Delta\theta = \pi/4$, $T = 0.5$ and $v_0 = 0.31$. The shape parameter for the cells $p_0 > 3.82$.

**Movie-16** Pattern obtained using the model with parameter $R_N = R_D = \rho = \mu = 1$, $\Lambda = -13.77$, $D_r = 0.1$, $v_0 = 3.1 \times 10^{-4}$, $\frac{\beta_j}{\beta_p} = 0.01$, $T = 0.94$ and $\Delta\theta = \pi/4$ and $\xi/D_r = 0$.

**Movie-17** Pattern obtained using the model with parameter $R_N = R_D = \rho = \mu = 1$, $\Lambda = -13.77$, $D_r = 0.1$, $v_0 = 3.1 \times 10^{-4}$, $\frac{\beta_j}{\beta_p} = 0.01$, $T = 0.94$ and $\Delta\theta = \pi/4$ and $\xi/D_r = 0.5$.

**Movie-18** Pattern obtained using the model with parameter $R_N = R_D = \rho = \mu = 1$, $\Lambda = -13.77$, $D_r = 0.1$, $v_0 = 3.1 \times 10^{-4}$, $\frac{\beta_j}{\beta_p} = 0.01$, $T = 0.94$ and $\Delta\theta = \pi/4$ and $\xi/D_r = 1.0$.

**Movie-19** Pattern obtained using the model with parameter $R_N = R_D = \rho = \mu = 1$, $\Lambda = -13.77$, $D_r = 0.1$, $v_0 = 3.1 \times 10^{-4}$, $\frac{\beta_j}{\beta_p} = 0.01$, $T = 0.94$ and $\Delta\theta = \pi/4$ and $\xi/D_r = 1.5$.

**Movie-20** Pattern obtained using the model with parameter $R_N = R_D = \rho = \mu = 1$, $\Lambda = -13.77$, $D_r = 0.1$, $v_0 = 3.1 \times 10^{-4}$, $\frac{\beta_j}{\beta_p} = 0.01$, $T = 0.94$ and $\Delta\theta = \pi/4$ and $\xi/D_r = 2.0$.

**Movie-21** Pattern obtained using the model with parameter $R_N = R_D = \rho = \mu = 1$, $\Lambda = -13.77$, $D_r = 0.1$, $v_0 = 3.1 \times 10^{-4}$, $\frac{\beta_j}{\beta_p} = 0.01$, $T = 0.94$ and $\Delta\theta = \pi/4$ and $\xi/D_r = 2.5$.

**Movie-22** Pattern obtained using the model with parameter $R_N = R_D = \rho = \mu = 1$, $\Lambda = -13.77$, $D_r = 0.1$, $v_0 = 3.1 \times 10^{-4}$, $\frac{\beta_j}{\beta_p} = 0.01$, $T = 0.94$ and $\Delta\theta = \pi/4$ and $\xi/D_r = 10.0$.

**Movie-23** Pattern obtained using the model for $R_N = R_D = \rho = \mu = 1$, $D_r = 0$, $\frac{\beta_j}{\beta_p} = 10^{-2}$, $\Delta\theta = \pi/4$, $v_0 = 3.1 \times 10^{-4}$, and $\Lambda = -13.77$ and $T = 0.6$.

**Movie-24** Pattern obtained using the model for $R_N = R_D = \rho = \mu = 1$, $D_r = 1$, $\frac{\beta_j}{\beta_p} = 10^{-2}$, $\Delta\theta = \pi/4$, $v_0 = 3.1 \times 10^{-4}$, and $\Lambda = -13.77$ and $T = 0.6$.

---

[1] G. R. Mirams, C. J. Arthurs, M. O. Bernabeu, R. Bordas, J. Cooper, A. Corrias, Y. Davit, S.-J. Dunn, A. G. Fletcher, D. G. Harvey, et al., Chaste: an open source c++ library for computational physiology and biology, PLoS Comput Biol **9**, e1002970 (2013).

[2] M. Ester, H. P. Kriegel, J. Sander, and X. Xu, Proceedings of the second international conference on knowledge discovery and data mining (kdd-96) (1996).
